# The domain architecture of JBP1 suggests synergy between J-base DNA binding and thymidine hydroxylase activity

**DOI:** 10.1101/502252

**Authors:** Athanassios Adamopoulos, Tatjana Heidebrecht, Jeroen Roosendaal, Wouter G. Touw, Isabelle Q. Phan, Jos Beijnen, Anastassis Perrakis

## Abstract

JBP1 (J-DNA Binding Protein 1) contributes to biosynthesis and maintenance of base J (β-D-glucosyl-hydroxymethyluracil), a modification of thymidine (T) confined to pathogenic protozoa. JBP1 has two known functional domains: an N-terminal thymidine hydroxylase (TH) homologous to the 5-methylcytosine hydroxylase domain in TET proteins; and a J-DNA binding domain (JDBD) that resides in the middle of JBP1. Here we show that removing JDBD from JBP1 results in a soluble protein (Δ-JDBD) with the N- and C-terminal regions tightly associated together in a well-ordered domain. This Δ-JDBD domain retains thymidine hydroxylation activity *in vitro*, but displays a fifteen-fold lower apparent rate of hydroxylation compared to JBP1. Small Angle X-ray Scattering (SAXS) experiments on JBP1 and JDBD in the presence and absence of J-DNA, and on Δ-JDBD, allowed us to generate low-resolution three-dimensional models. We conclude that Δ-JDBD, and not the N-terminal region of JBP1 alone, is a distinct folding unit. Our SAXS-based model supports the notion that binding of JDBD specifically to J-DNA can facilitate hydroxylation a T 12–14 bp downstream on the complementary strand of the J-recognition site. We postulate that insertion of the JDBD module in the Δ-JDBD scaffold during evolution provided a mechanism to synergize between J recognition and T hydroxylation, ensuring inheritance of J in specific sequence patterns following DNA replication.

## INTRODUCTION

Base J (β-D-glucosyl-hydroxymethyluracil) is a modified base that replaces 1% of thymine in kinetoplastid flagellates^1^, such as *Trypanosoma, Leishmania* and *Crithidia*. In *Leishmania*, 99% of J is located in telomeric repeats^2–4^ whereas 1% is in internal chromosomal positions (iJ) in positions where transcription starts^5^ or stops^6^.

Biosynthesis of J occurs in two-steps. First, the 5-methyl group of specific T’s in the genome is 0hydroxylated, forming hydroxymethyluracil (hmU). Second, a glucose molecule is transferred to hmU, resulting in J^7–9^. The first step is catalyzed by both J-DNA Binding Proteins 1 and 2 (JBP1 and JBP2), that have a distinct thymidine hydroxylase domain (TH) in their N-terminus^10^. The hydroxylation of T in oligonucleotides is dependent on the presence of Fe(II), 2-oxoglutarate (2OG)^11^. The discovery of the hydroxylation function of JBP1 has led to the discovery of the function of the mammalian TET family of enzymes that convert 5-methylcytosine (5mC) to 5-hydroxymethylcytosine (hmC)^12^, and have crucial roles in epigenetic regulation through modification of 5mC to hmC. Several structures of TET and TET-like hydroxylase domains have been determined^13–15^, also in complex with 5mC and hmC, providing significant insight in 5mC hydroxylation. However the limited sequence similarity between TET and JBP, underlined by large deletions and insertions in the TH fold, makes it impossible to deduce the structures of JBP1 and JBP2 from that of TET.

JBP1, but not JBP2, specifically recognises base J in DNA^16^. This recognition is mediated by a short ~150 residue domain in the middle of JBP1, the J-DNA binding domain (JDBD)^17^. JDBD recognises J-DNA with high affinity (~10 nM) and remarkable specificity over normal DNA (~10,000 fold). The structure of JDBD revealed a novel variant of the helix-turn-helix domain, with an unusually elongated turn between the recognition and the supporting helix. Importantly, we have shown that a single residue (Asp-525) in the recognition helix is almost entirely responsible for the specificity towards J-DNA, as the D525A mutation abrogated specificity towards J-DNA both in vitro and in vivo.

JBP1 recognizes and binds J-DNA in two steps^18^. Pre-steady state kinetic data revealed that the initial binding of JBP1 to glucosylated DNA is very fast and followed by a second, much slower and concentration independent step. From this observation and small-angle neutron scattering (SANS) experiments, we inferred that JBP1 undergoes a conformational change upon binding to DNA, and postulated that this may allow the hydroxylase domain of JBP1 to make contact with the DNA and hydroxylate Ts in spatial proximity.

From what we know about the mechanism of J biosynthesis, it follows that the highly restricted distribution of J-base must be co-determined by the thymidine hydroxylases (JBP1 and JBP2) that catalyze the initial step in J synthesis. Using SMRT sequencing of DNA segments inserted into plasmids grown in *Leishmania*^19^, it has been shown that J modification usually occurs near G-rich sequences potentially capable of forming G-quadruplexes and at pairs of Ts on opposite DNA strands, separated by 12 nucleotides. That led Genest *et al*^19^ to propose a model in which JBP2 is responsible for initial J synthesis; then JBP1 binds to pre-existing J and hydroxylates another T that typically resides 13 bp downstream (but not upstream) on the complementary DNA strand. This model provides a mechanism explaining how JBP1 can maintain existing J following DNA replication.

On the basis of these results we developed the conformational change model presented in ref. 18: We postulated that J-binding docks JBDB on duplex DNA, allowing the TH domain to come into contact with a T that is preferably 13 bp downstream the complementary strand and hydroxylate it. To further test this model, we set out to understand the domain organization of JBP1 and describe their three-dimensional organization alone and in relation with DNA.

Here, we present new deletion mutants and hydroxylase activity data of JBP1, that result to a new definition for the TH domain, and we also study the domain architecture of JBP1 by small angle X-ray scattering (SAXS). Developments in SAXS, namely the improved software and hardware, have established SAXS as a powerful tool for the analysis of molecular structures, also in the case of multi-domain, flexible molecules^20–24^. Coupling size-exclusion chromatography to SAXS (SEC-SAXS) allows separating complexes from the constituent partners, degradation products, and eventual contaminants, allowing the determination of particle size and shape of macromolecules. Collecting and analysing SAXS data on different JBP1 deletion mutants and their complexes with J-DNA, allowed us to generate low-resolution three-dimensional models of JBP1 and its complex with DNA. Our data suggest synergy between the TH and JDBD domain (that is likely a recurrent fold in nature, not solely confined to JBP1 orthologues), is an evolutionary adaptation, crucial for replicating epigenomic information in kinetoplastids.

## RESULTS

### The N- and C-terminal regions of JBP1 behave as a single folding unit

Countless previous attempts to truncate JBP1 either C-terminally or N-terminally (before or after the JDBD), to obtain the N-terminal TH domain or a putative C-terminal domain, had invariably failed to yield soluble protein in our hands. The crystal structure of JDBD showed that the N- and C-termini of this domain are in proximity (Figure 1A). As JDBD is in the middle of the JBP1 sequence (Figure 1B) we decided to test the possibility that the JDBD is an insertion domain into a “TH domain” fold that spans the rest of the JBP1 sequence. To validate this hypothesis, we replaced the JDBD domain with a connecting linker. Remarkably, the resulting protein (JBP1^1-382/561–882^, Δ-JDBD) expressed well in soluble form, and could be purified in good amounts (Figure 1C). Intrigued by that, we wanted to examine if the N-terminal (1–382) and the C-terminal (561–882) regions form a single folding unit, or behave as separate domains. We therefore introduced a 3C protease cleavage site in the connecting linker between the two halves of Δ-JDBD (Figure-1B). Overexpression of this Δ-JDBD-3C construct also yielded soluble protein (Figure 1C). Incubation with 3C protease over a period of 13–36hr resulted in the protein chain to be cleaved in two, and these bands could be observed on an SDS-PAGE (Figure 1C). However, when we run the cleaved protein on a size exclusion chromatography (SEC) column in the absence of detergent, the elution profile showed a single symmetric peak (Figure 1D) of approximately the same molecular weight as Δ-JDBD, while the SDS-PAGE analysis of the eluted fractions confirmed that this peak has both bands present. This strongly suggests that the two polypeptides behave as one protein, indicating a strong interaction between the two termini. This indicates that both termini should be part of the same folding unit, which is the folding unit necessary to provide a functional catalytic site containing the TH activity. This experiment led us to revisit our previous view of JBP1, with an N-terminal TH domain, followed by the JDBD domain and a mysterious C-terminal region. We now hypothesized, that two folding units compose JBP1: the Δ-JDBD and the JDBD that is “inserted” in the Δ-DJBD folding scaffold.

**Figure 1:**
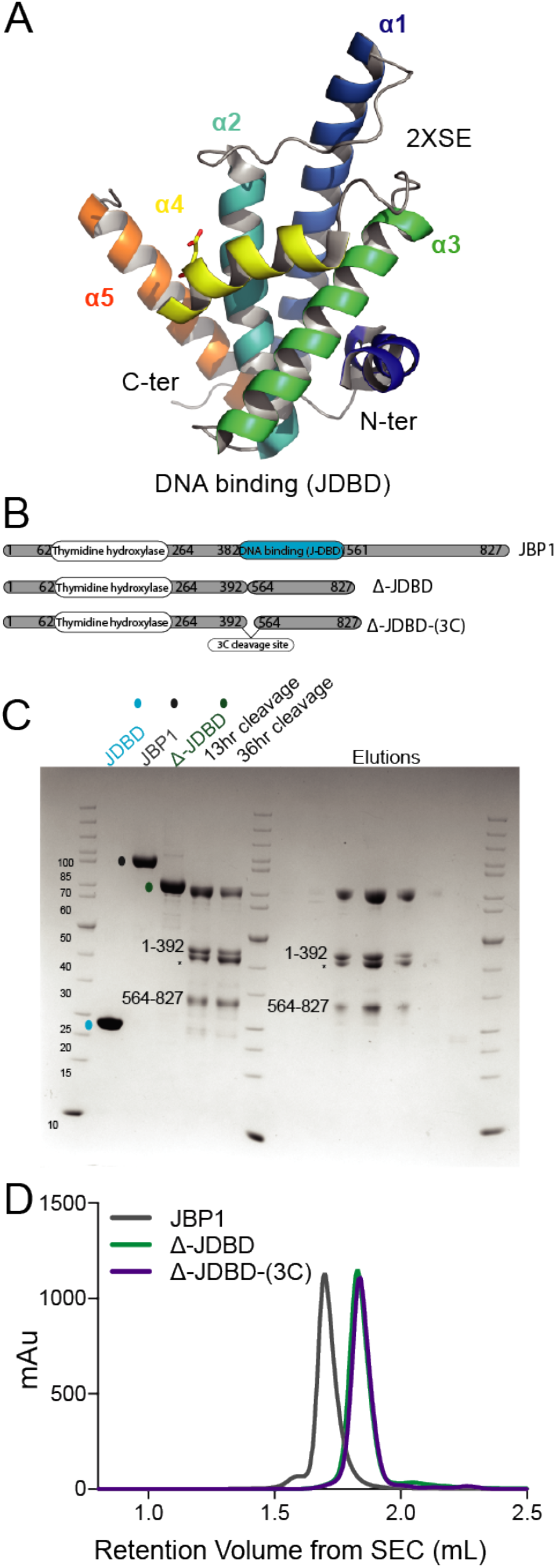
Domain organization of JBP1. **(A)** cartoon representation of JDBD crystal structure, showing the characteristic helical bouquet fold containing the helix-turn-helix motif. Asp-525 is shown as stick. **(B)** Constructs used in this study; full-length JBP1, Δ-JDBD-3C, Δ-JDBD. **(C)** SDS-PAGE showing the overexpressed constructs in the following order: Marker, JDBD, JBP1, Δ-JDBD-3C, Δ-JDBD-3C products after 13 and 36 hr incubation with 3C protease, marker, SEC elution fractions of Δ-JDBD-3C, after incubation with 3C protease. **(D)** Chromatogram of the SEC for JBP1, Δ-JBP1 and Δ-JDBD-3C; Δ-JBP1 before and after cleavage with 3C protease are identical. The N- and C-terminal regions of Δ-JDBD-3C, elute in the same peak (see panel (C) above, suggesting that there is a strong interaction between them.

### JBP1, JDBD, Δ-JDBD are all well-folded globular domains in solution

To further characterize Δ-DJBD in solution, we performed Small Angle X-ray Scattering experiments on JBP1, JDBD, and Δ-JDBD. All samples were injected in a size exclusion chromatography column (SEC), and the SAXS profile (as well and the absorption spectrum) were analysed under flow. All samples eluted as single peaks from the SEC and the SAXS curves from the absorption peak region were averaged to obtain a scattering curve for each component (Figure 2A and Supplemental Table 1). Standard analysis tools from the ATSAS^32^ and SCÅTTER^20^ suites were used to obtain model-independent parameters (Table 1). To confirm that all samples were properly folded we performed a dimensionless Kratky plot analysis (Figure 2B). This plot, allows to compare the shape of particles independent of their size, and shows that both JBP1 and Δ-JDBD have a similar profile with a maximum close to 1.104, characteristic of compacted and folded molecules. The JDBD peak is shifted slightly right and upward, suggesting that JDBD is less globular but compacted (which agrees with the crystal structure shape). The pair distribution function (Figure 2C) is compatible with this analysis, and together they confirm our interpretation of the biochemical experiments, suggesting that Δ-JDBD is a stable single domain.

**Figure 2:**
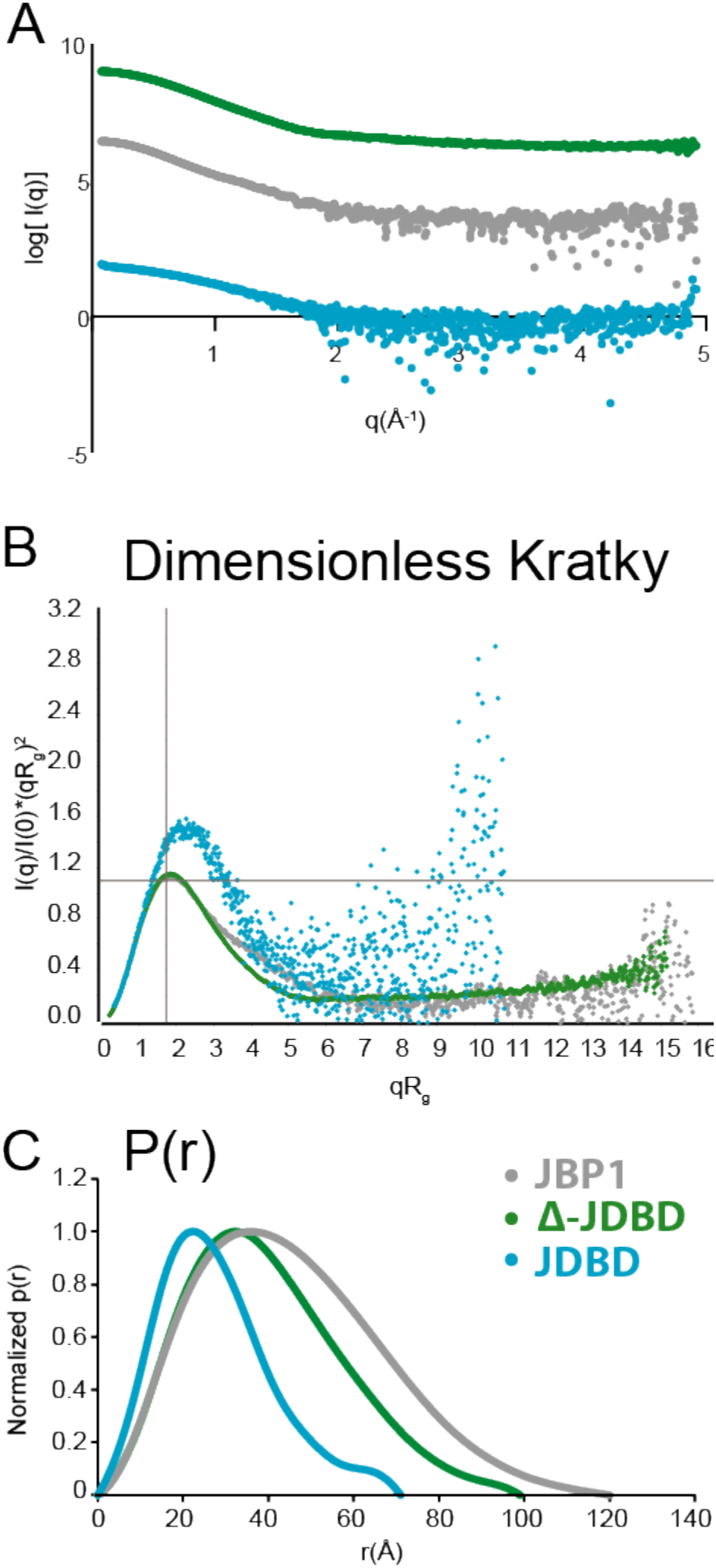
Folding of JBP1, Δ-JDBD, JDBD and 23 J-DNA. **(A)** Experimental scattering curves of all components. **(B)** Dimensionless Kratky plot; the 1.104 maximum for an ideal compacted molecule is shown as the intersection between the 2 grey lines; **(C)** normalized pair distribution function for all components.

**Table 1.** Model independent parameters for all samples used in this study.

| | $R_{g\text{-Guinier}} (\text{\AA})$ | $R_{g\text{-reciprocal}} (\text{\AA})$ | $D_{\text{max}} (\text{\AA})$ | $MW_{\text{theory}} (\text{D})$ | $V_{\text{Porod}}$ | $V_c$ |
| --- | --- | --- | --- | --- | --- | --- |
| **Monomers** |  |  |  |  |  |  |
| JBP1 | 34.3 | 34.3 | 120 | 93403 | 98.7 | 610.1 |
| Δ-JDBD | 30.6 | 30.6 | 99 | 72118 | 76.4 | 505.5 |
| JDBD | 24.8 | 24.8 | 71 | 21285 | 28.3 | 260.9 |
| J-23-DNA | 22.5 | 22.5 | 73 | 14241 | 16.2 | 201.7 |
| J-15-DNA | 15.8 | 15.8 | 51 | 9301 | 10.3 | 131.8 |
| **Complexes** |  |  |  |  |  |  |
| JBP1:J-23-DNA | 40.98 | 40.98 | 141 | 107644 | 107.1 | 685.8 |
| JBP1:J-15-DNA | 35.6 | 36.29 | 120 | 102704 | 104 | 641.9 |
| JDBD:J-23-DNA | 21.7 | 21.67 | 70 | 35526 | 34.73 | 241.1 |

### Δ-JDBD is a catalytically active domain that has thymidine hydroxylase activity

We first established a mass spectrometry-based assay to measure TH activity of JBP1 *in vitro*. We used purified proteins with a 14-mer oligonucleotide in conditions similar to those reported in ^46^ to convert T to hmU, which was measured by quantitative liquid chromatography mass spectrometry, after converting the oligonucleotide to nucleosides (see Methods for details). This activity was fully dependent on the presence of the co-factor 2-oxoglutarate, and showed a modest but appreciable dependency to Fe^+2^, ascorbic acid as a reducing agent, and to buffer degassing (Supplementary Figure 1). We then monitored the rate of catalysis over time for both JBP1 and Δ-JDBD. Δ-JDBD was clearly active, but showed a catalytic rate of about 17-times lower than wild type JBP1 (Figure 3).

**Figure 3:**
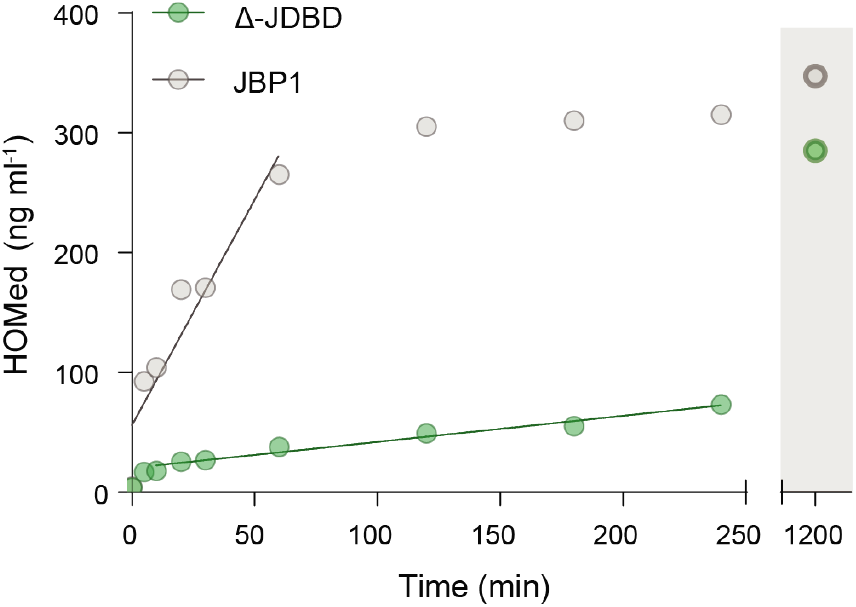
The apparent catalytic rates of JBP1 and Δ-JDBD. The linear part of the reaction curve (5–60 min for JBP1, 10–240 minutes for Δ-JDBD) are fitted with a line to estimate the rates.

These experiments clearly establish that Δ-JDBD is a well-folded active TH domain, which is not disrupted by splicing out the JDBD domain. We thus decided to then characterise the relative domain architecture between Δ-JDBD and JDBD in the context of the JBP1 protein.

### Modelling of JBP1 as a two-domain (Δ-JDBD and JDBD) molecule shows flexibility for JDBD

First, we compared further JBP1 to Δ-JDBD by examining the Porod-Debye plot (Figure 4A). The presence of a plateau for Δ-JDBD indicates that it forms a distinct particle with sharp scattering contrast. This feature is not present in JBP1, indicating a more diffuse scattering contrast for the full length JBP1. The same is observed calculating the packing densities for both Δ-JDBD and JBP1: Δ-JDBD has a packing density of 0.91 g cm^−3^ compared with 0.79 g cm^−3^ of JBP1. The observed diffusion in scattering contrast and the reduced packing density in the wild type, full-length, JBP1 compared to Δ-JBP1 suggest that the JDBD domain is flexible with respect to the Δ-JDBD scaffold.

**Figure 4:**
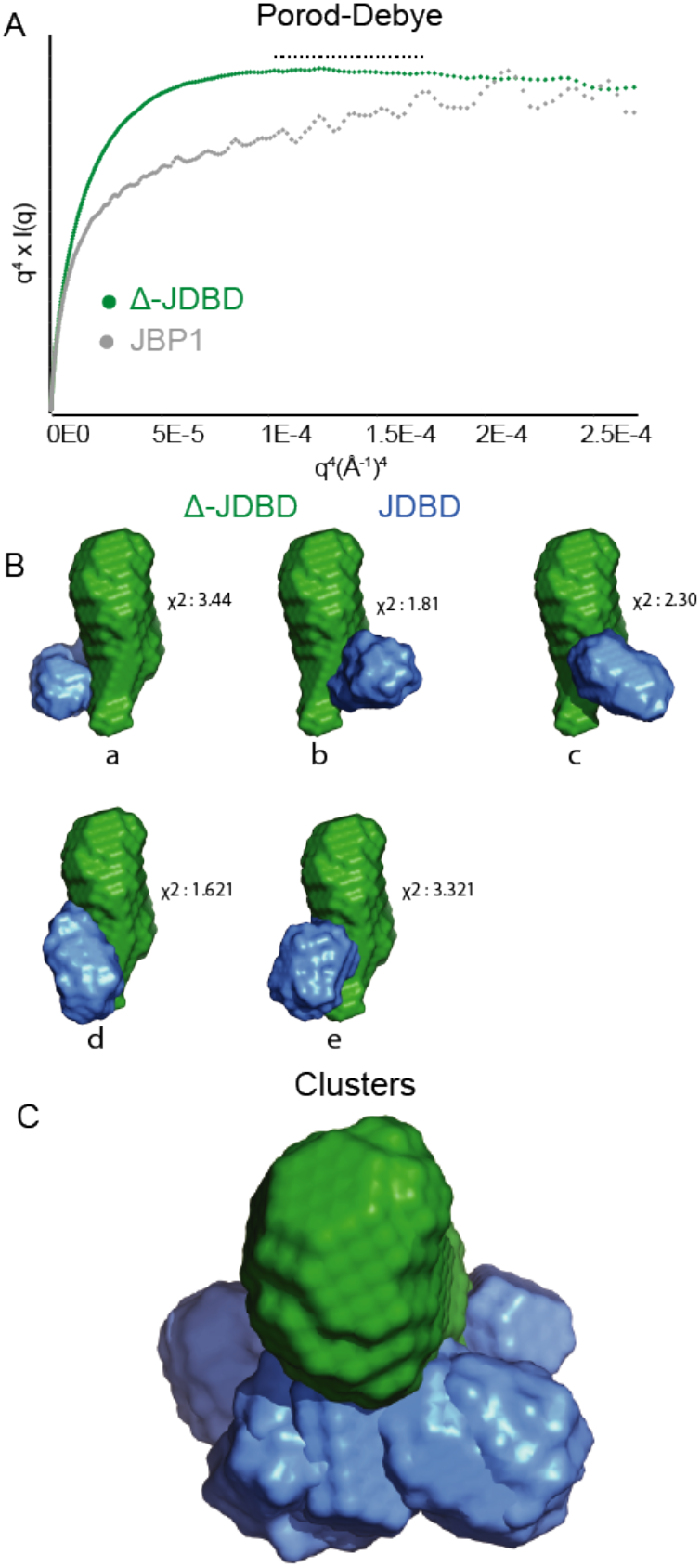
MONSA modelling of JBP1. **(A)** Porod-Debye plot demonstrating a loss of the plateau when JDBD is present, suggesting that JBP1 is more flexible. **(B)** Clustered MONSA models of JBP1; Δ-JDBD and JDBD are shown in green and blue, respectively; a total five clusters were identified and the fitting to the experimental data of the calculated intensities for each model is shown based on χ^2^. **(C)** A representation of all possible JDBD orientations shown along the long axis of Δ-JDBD. Figures were prepared using the program ScÅtter and Pymol.

To validate this hypothesis we decided to create *ab initio* three-dimensional models based on the SAXS data. As we have SAXS data for Δ-JDBD and JDBD alone, as well as for both of them together (JBP1), we decided it is more appropriate to model them with the procedures developed for macromolecular complexes and multi-domain proteins^32,35^. The program MONSA from the ATSAS suite^32^ seeks to identify so called “multi-phase” models (each “phase” being a rigid domain) that fit simultaneously the scattering data describing each phase (domain) separately, and their complex. We defined two phases, Δ-JDBD and JDBD, which make up a “complex”, JBP1. Twenty models were created by MONSA to fit the three available scattering data sets. Details for this and subsequent MONSA modelling runs are in Table 2.

**Table 2.** Details for the phases used in MONSA for modeling, and all resulting clusters, for all complexes and sub-complexes used in this study.

| Complex | Phase 1 | Phase 2 | Cluster-1 | Cluster-2 | Cluster-3 | Cluster-4 | Cluster-5 |
| --- | --- | --- | --- | --- | --- | --- | --- |
| JBP1 | Δ-JDBD | JDBD | 7 | 3 | 2 | 2 | 2 |
| JBP1:23-J-DNA (I) | JBP1 | 23-J-DNA | 7 | 5 | 4 | 3 | - |
| JBP1:23-J-DNA (II) | Δ-JDBD | JDBD:23-J-DNA | 10 | 6 | 2 | - | - |
| JBP1:15JDNA | JBP1 | 15-J-DNA | 14 | 3 | 2 | - | - |

An examination of the individual models showed that the JDBD domain adopts multiple conformations with respect to the Δ-JDBD scaffold. Clustering analysis with the program DAMCLUST from the ATSAS suite^32^ identified five clusters (Figure 4B), all of which have JDBD located on the same end of the elongated Δ-JDBD scaffold, but in various positions around the long axis of the Δ-JDBD domain (Figure 4C). If Δ-JDBD is viewed as an ellipsoid, the JDBD is consistently positioned towards one half of the ellipsoid, but adopts multiple conformations around the long axis of the ellipsoid. This analysis is compatible with the model-independent analysis of the SAXS data and strengthens our previous hypothesis, that the JDBD domain is flexible with respect to the Δ-JDBD scaffold.

### Binding of JBP1 to J-DNA leads to reduced flexibility of the JDBD domain

We have previously shown that JBP1 and J-DNA complex formation is accompanied by a conformational change^18^. Our new data allow us to formulate the hypothesis that this conformational change might be the ordering of the JDBD: when JBP1 binds to DNA, JDBD might adopt a more defined conformation with respect to the Δ-JBP1 scaffold. To test this hypothesis, we used the SEC-SAXS data on JBP1 in complex with 23-mer J-DNA (J-23-DNA). SEC-SAXS data were first collected for J-23-DNA, which had the expected parameters for an elongated molecule (Table 1 and Supplemental Table 1). The complex between J-23-DNA and JBP1 eluted from the SAXS column as a single peak, and was confirmed by the 280/260 nm absorption ratio; the averaged SAXS profile from the elution peak are shown in Figure 4A; model-independent parameters are in Table 1 and SEC details in Supplemental Table 1.

While visual inspection of the scattering intensity for JBP1 alone and the complex suggests that they are very similar (Figure 5A), plotting the intensity ratio of the two datasets shows that the molecular form factors for the two datasets have prominent differences, as we observe strong features throughout the curve (Figure 5B). Analysis of the data also shows that an increase in R_g_ was accompanied by an increase in D_max_, suggesting that J-DNA binds away from the centre of mass (Table 1). The volume-of-correlation, V_C_, is higher in the J-DNA-bound state, similar to the Porod volume that also increases by 14,000 Å^3^ in the presence of J-23-DNA. Finally, examination of the dimensionless Kratky plot reveals a shift away from the Guinier-Kratky point (1.104) indicating that upon J-DNA binding, JBP1 has a more elongated shape (Figure 5C). All these data establish that the complex of JBP1 and J-23-DNA is formed, and that the J-23-DNA binds away from the JBP1 centre-of-mass resulting in a more elongated particle.

**Figure 5:**
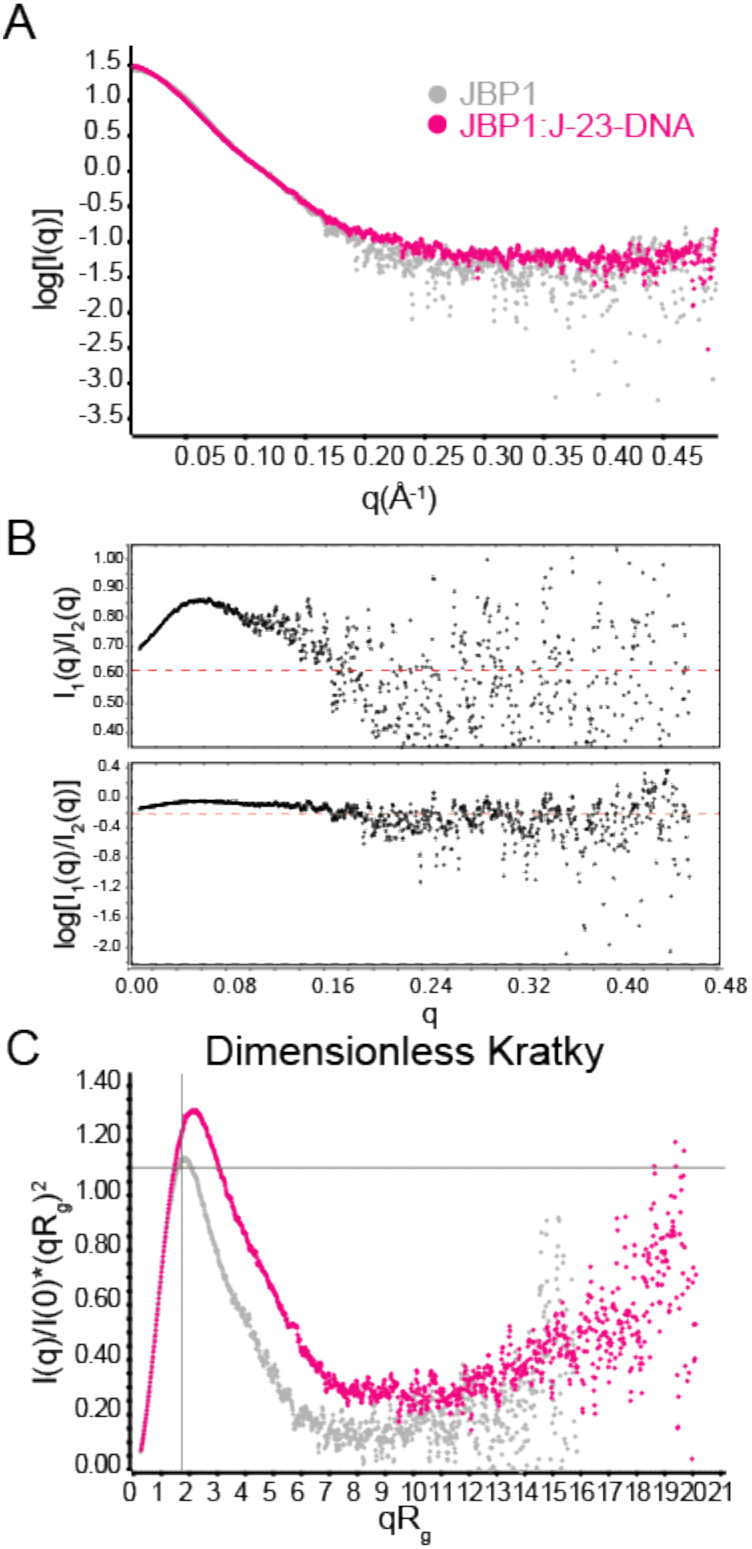
Comparison of JBP1, in presence and absence of J-23-DNA. **(A)** Experimental scattering curves of JBP1 and JBP1:J-23-DNA. **(B)** Ratio plots of the two datasets. **(C)** Dimensionless Kratky plot for both datasets.

To visualize the relative position of J-DNA in the complex, we again used the program MONSA. We defined two phases, J-23-DNA and JBP1, and calculated 20 models consistent with the scattering data for JBP1, J-23-DNA and the JBP1:J-23-DNA complex. Cluster analysis with DAMCLUST, resulted in four major clusters (Figure 6). In all clusters J-DNA is located towards one end of the JBP1 ellipsoid, away from its centre of mass, compatible with the positioning of the JDBD domain. In contrast with the two-body modelling of JDBD and Δ-JBP1, the clusters are rather similar, suggesting that J-23-DNA binds in similar conformations. As previous SAXS data on the complex between JDBD and J-DNA suggested that this is a rigid complex without conformational flexibility^25^, this postulates that this JDBD: J-23-DNA rigid complex is now in one conformation with respect to Δ-JDBD. In other words, this analysis is compatible with the hypothesis that the JDBD gets ordered upon J-DNA binding. To confirm these finding, we repeated the procedure with 15mer J-DNA (J-15-DNA); the results (Table 1 and Supplementary Figure 2) lead to the same conclusions.

**Figure 6:**
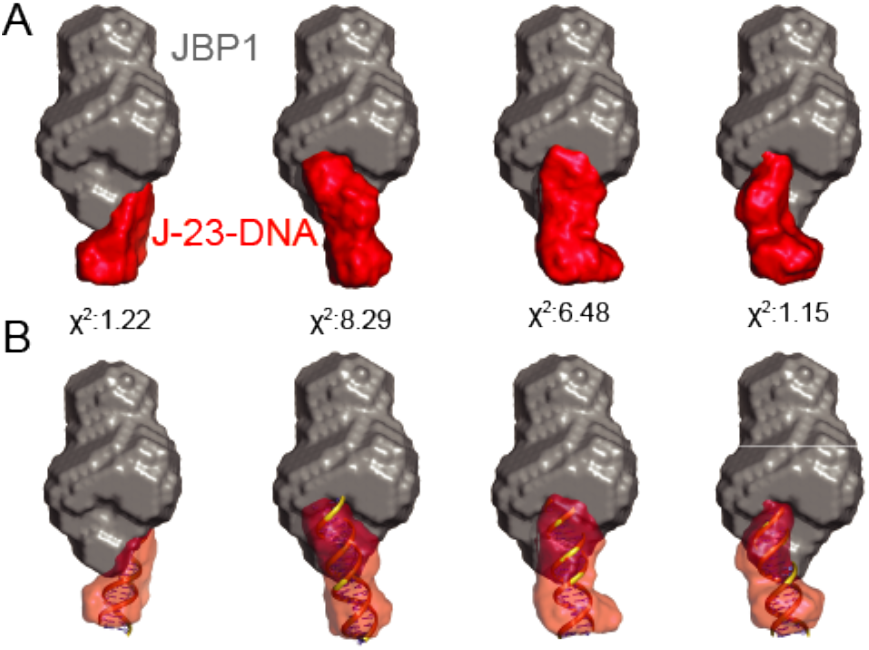
Model of JBP1:J-23-DNA. **(A)** Clusters of all JBP1:J-23-DNA MONSA models, with the χ^2^ fitting of the calculated intensities to the experimental data. **(B)** The J-23-DNA is superimposed on the molecular envelope generated by MONSA. Superposition was performed using SUPCOMB from the ATSAS suite. Figures generated using Pymol.

We then proceeded to test our hypothesis for the ordering of the JDBD upon complex formation, using a different modelling approach. For this, we treated the JDBD complex with J-DNA as a rigid body ^25^ and defined two different phases: Δ-JBP1 and the JDBD:J-23-DNA complex. We again used MONSA for calculating 20 models consistent with the scattering data for Δ-JBP1, and the JDBD:J-23-DNA and JBP1:J-23-DNA complexes. Cluster analysis with DAMCLUST resulted in three clusters (Supplementary Figure 3). Consistent with the previous analysis, these clusters are fairly similar, confirming the reduced flexibility of the complex of JBP1 with J:DNA, and again show the DNA and JDBD located towards one end of the complex.

As we have an atomic model for the JDBD:J-23-DNA complex^25^, we were able to place it in the respective dummy atom model using SUPCOMB^39^. Thus, we created a pseudo-atomic hybrid model, where the Δ-JDBD (for which we do not have an atomic model) is shown as the dummy atom model, and JDBD and J-23-DNA are all-atom models. Using CRYSOL^42^ we evaluated the fit of both the dummy atom reconstruction and the pseudo-atomic hybrid model against the SAXS curve for JBP1:J-23-DNA. The dummy atom reconstructions for the two most populated clusters, with ten and six members respectively, show the best fit to the experimental data (χ=2.32 and χ=1.68 respectively). However, the pseudo-atomic hybrid model corresponding to the most populous cluster shows a considerably better fit to the experimental data (χ=6.87) compared to the second cluster (χ=38.03). Thus, we consider this model as the most likely interpretation of our experimental data for the complex of full length JBP1 with J-DNA (Figure 7).

**Figure 7:**
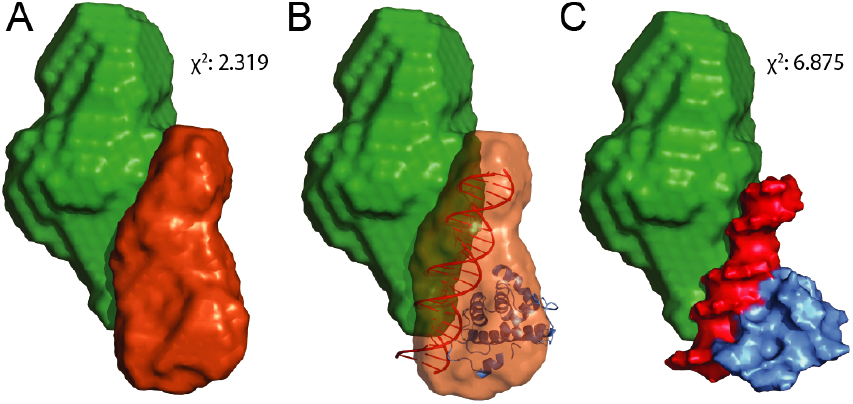
Model of JBP1 in presence of JDNA. **(A)** Clustered MONSA model showing the relative positions of Δ-JDBD and JDBDJ-23-DNA sub-complex. **(B)** A pseudo-atomic model of JDBD:J-23-DNA fitted in the molecular envelope of the generated model. **(C)** Surface representation of all components. Figures were made using Pymol.

This pseudo-atomic model now shows the position of the J-base and the most likely orientation of the DNA. Interestingly, in this model, the T base 13bp away from J in the complementary strand gets in contact with the Δ-JDBD domain that contains the TH activity.

## DISCUSSION

The discovery that JBP1 has an N-terminal TH domain sequence signature, which likely functions as a thymidine hydroxylase^10^, was an important finding for the field of J biosynthesis. Perhaps more remarkably, this sparked a revolution for the methylcytosine to hydroxymethylcytosine field^12^. Together with subsequent experimental proof of the TH activity hypothesis^11^, these findings established the notion that JBP1 consists of an N-terminal TH domain, followed by a J-DNA binding domain^25^, and a C-terminal sequence which received little attention. Here, we show that the JDBD should be seen as an insertion domain within a single TH domain that spans the N-terminal and C-terminal sequence regions of JBP1, and behaves as a single folding unit in solution; Δ-JDBD. We note that a JBP1 construct spanning residues 1–451 which has been previously reported to have hydroxylase activity^47^, as well as numerous other constructs of the N-terminal from a variety of species, does not yield soluble protein in our hands. The sole exception to that rule has been the Δ-JDBD domain. Remarkably, this new folding unit is functional as a thymidine hydroxylase, in a new enzymatic activity assay that we developed, having an apparent catalytic rate about 17 times slower compared to full-length JBP1. We suggest that this lower rate is explained by the inability of Δ-JDBD to bind to DNA, bringing it in proximity to its T base substrates.

We have previously shown that binding of JBP1 to J-DNA is followed by a conformational change of JBP1^18^. Here, we extend this model, providing data that this conformational change represents a transition of JDBD: while JDBD is flexible with respect to the Δ-JDBD scaffold in the absence of J-DNA, it gets ordered in the presence of J-DNA. This is in agreement with our previous observation from SANS data^18^ that the protein apparent radius of gyration (R_g_) is reduced upon complexation with J-DNA. This is consistent with JDBD sampling a more defined conformation space and reducing the apparent size of the protein particle.

The complex between JDBD and J-DNA is a well-defined rigid structure, as shown by our previous structural analysis^44^ and current data. Our current analysis of Δ-JDBD and the JBP1 complex with J-DNA shows these to be also fairly rigid. These allowed us to propose a pseudo-atomic hybrid model, showing the orientation of J-DNA with respect to the Δ-JDBD domain that contains the TH catalytic activity (Figure 7). In that model, the DNA gets in contact with the Δ-JDBD. The J-23-DNA sequence we used for these experiments contains both the J that is recognized by the JDBD, but also a complementary strand sequence that is amenable to hydroxylation. Interestingly, in our most probable model, the T that lies 13bp downstream in the complementary strand comes in close contact with the Δ-JDBD domain that has the TH activity. Thus, our structural analysis supports the hypothesis that the JDBD domain of JBP1 binds to J, and the Δ-JDBD domain then undergoes a conformational change allowing it to reach and hydroxylate a T 13 bp on the complementary strand. In this way JBP1 is able to maintain existing J following DNA replication (Figure 8).

**Figure 8:**
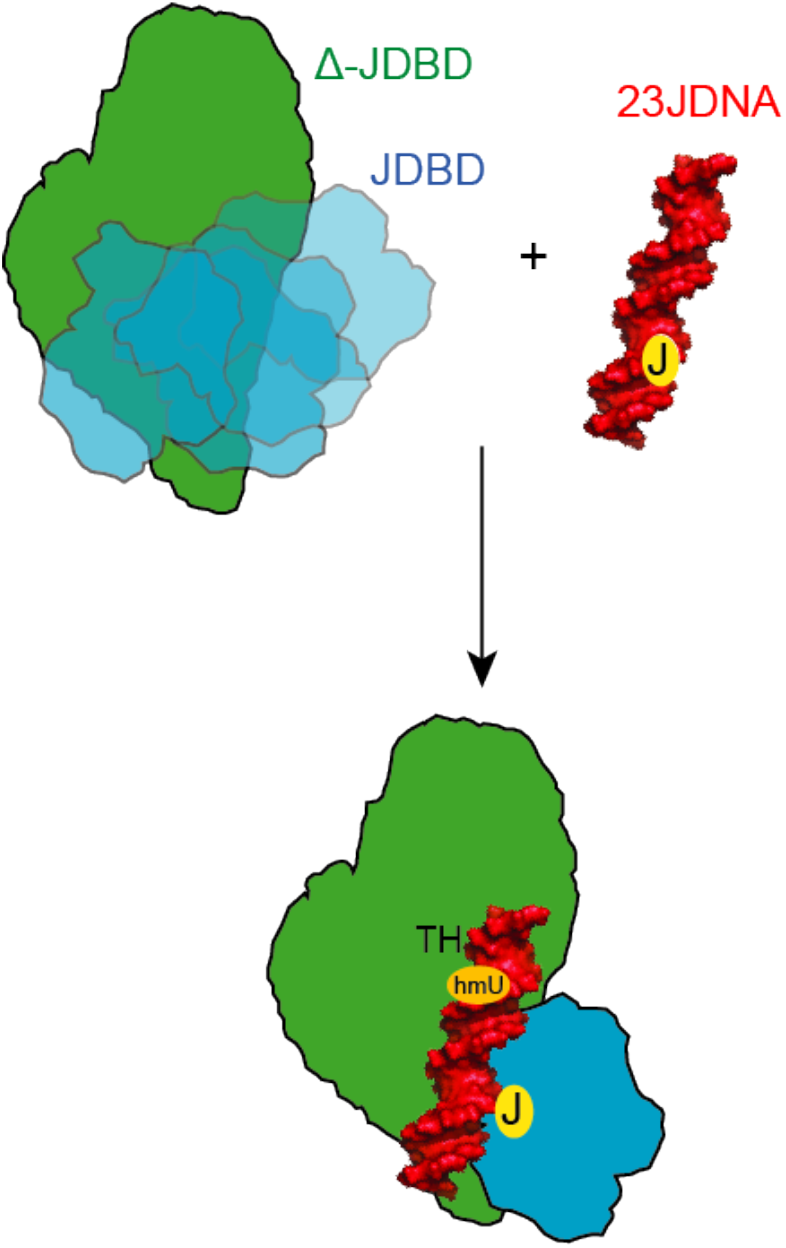
A model suggesting the mechanism for the recognition and maintenance of base-J. JDBD recognizes and binds base-J with high affinity, result in a conformational change that could stabilize the JDBD domain. Upon binding, JDNA is an orientation that allows the TH-domain to be in close proximity to the thymine base that is located 13 bp downstream on the complementary strand to promote its hydroxylation, and therefore maintenance of J at specific positions in the genome of kinetoplastids.

From an evolutionary perspective, it is reasonable to presume that the thymidine hydroxylation activity to make hydroxymethyluracil precedes the glucosylation step to make J. We hypothesize that the last evolutionary step was the acquisition of J-binding activity, to guide the TH activity to areas of pre-existing J to replicate that epigenetic marker in kinetoplastids. As we show here, JDBD has likely been acquired by JBP1 through an insertion event that did not disturb the TH scaffold. Based on these observations, one would expect to find JDBD homologues – with or without specificity for J-DNA binding – in additional proteins. As sequence searches in public databases do not reveal clear homologues of the JDBD outside the context of JBP1 orthologues, we performed structural similarity searches using DALI^43^ (see Methods for details). These searches revealed two new structural homologues additional to MogR^48^, which we have previously described^25^. The closest structural homologue of JDBD is AcrF3, belonging to a family of proteins produced by bacteriophages to inactivate the CRISPR–Cas bacterial immune system^45^; the other homologue is a C-Terminal helical domain (CHCT) of the chromatin remodelling protein CHD1^49^. While JDBD, MogR, and CHCT clearly have a conserved positive patch for interaction with DNA (Figure 9), this patch is absent in AcrF3. Interestingly, anti-CRISPR (Acr) proteins bind the Cas complexes blocking recognition of double-stranded DNA substrates^50,51^: speculating that the ancestry of Acr proteins is related to the JDBD, MogR and CHCT DNA recognition domains, this might present an extreme example of repurposing a DNA-recognition structural domain for preventing DNA-recognition, or vice versa. That data suggests that the JDBD DNA-recognition scaffold might be considerably more widespread and not confined to specific J recognition; as complete sequences of protozoan species will continue to be fully assembled, more JDBD-like domains will likely be identified, inside or outside the J-biosynthesis pathway.

**Figure 9:**
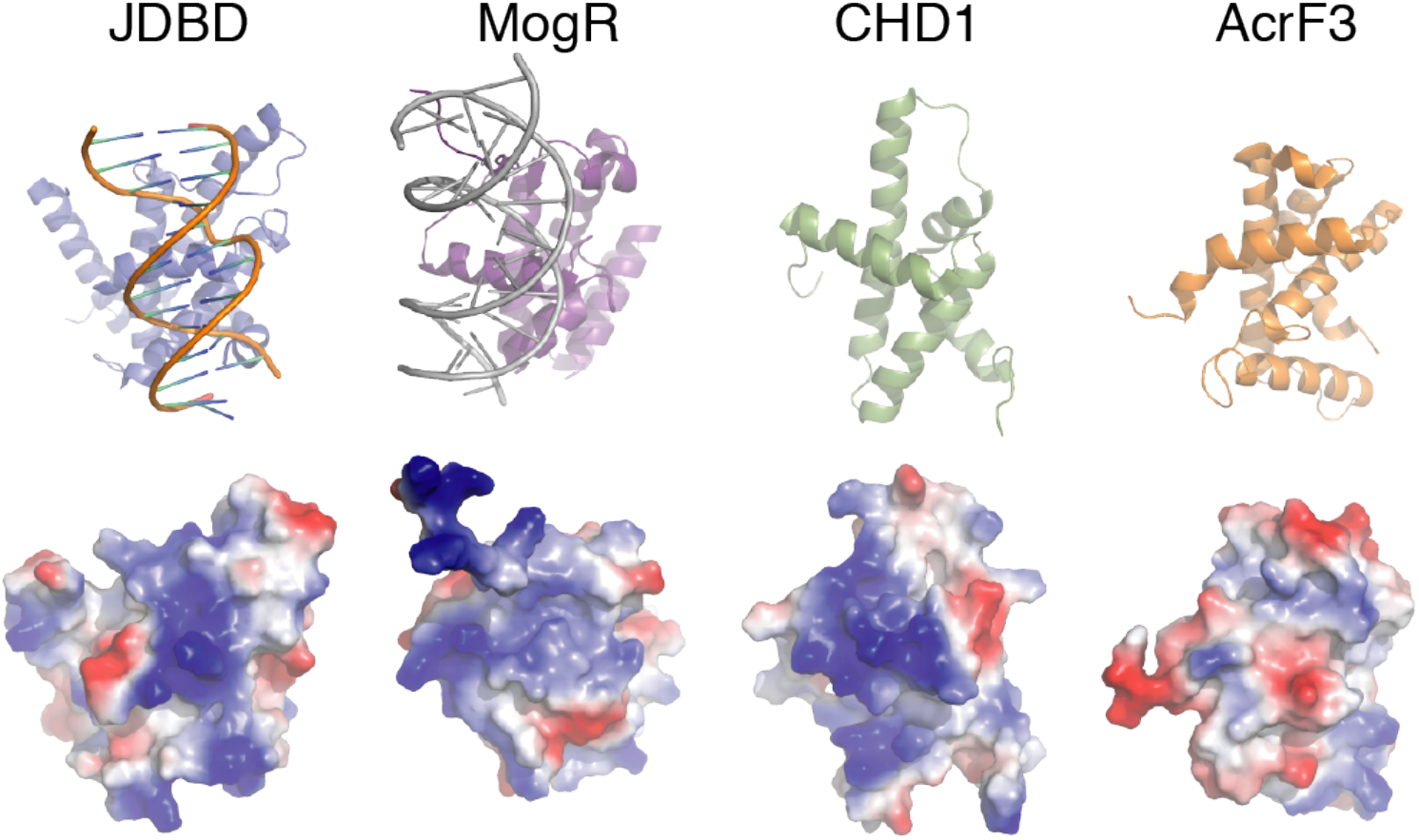
Cartoon models of the JDBD and homologues in the PDB (top) and the corresponding surface representations coloured by electrostatic potential. The positive blue patches are characteristic for the interaction with DNA as shown in the SAXS-based model of JDBD and the MogR crystal structure; this positive patch is entirely missing in the AcrF3 protein.

## Supplementary Data are at the end of the manuscript

## ACKNOWLEDGEMENT

We thank Piet Borst for extensive discussions and suggestions to improve the text of this manuscript and Piet Borst and Henri van Luenen for helping to establish the JBP1 activity assays.

## METHODS

### Cloning of Δ-JDBD JBP1 and Δ-JDBD-3C JBP1

A JBP1 synthetic gene encoding the sequence for *Leishmania tarentolae*^25^ was used as the template for all constructs in this study. Primers to delete the JDBD domain and to replace it with the 3C-protease DNA sequence were designed using the ProteinCCD software^26^. To create the Δ-JDBD construct, the JDBD domain was deleted using mutagenesis PCR and primers *Del_fw 5’- CTC GTC TGG GTG GTT TCT CTG AAA CCT CTC ACG AAA AAC GTG CTA ACT GGC TG −3’* and *Del_rev 5’- CAG CCA GTT AGC ACG TTT TTC GTG AGA GGT TTC AGA GAA ACC ACC CAG ACG AG −3’*. The generated construct contained the N-terminal part from residues 1–392 followed by the C-terminal part residues 564–827 without any connecting linker in between. To generate the Δ-JDBD-3C construct a 3C protease cleavage site was introduced using primers *Del-DB-3C_fw 5’-CTC GTC TGG GTG GTT TCT CTG AAA CCC* ***TGG AAG TGC TGT TTC AGG GCC CGT*** *CTC ACG AAA* −3’ and *Del-DB-3C_rev 5’-CAG TTA GCA CGT TTT TCG TGA GAC* ***GGG CCC TGA AAC AGC ACT TCC AG****G GTT TCA GAG AAA −3’* respectively (highlighted is the DNA sequence for the 3C-protease cleavage site).

### Expression and purification of recombinant proteins

All constructs of JBP1, JDBD and Δ-JDBD were inserted in the NKI-LIC-1.1 vector^27^ and produced as soluble proteins in *E.coli*. BL21 (DE3) T1R cells were used for protein overexpression. Protein production was induced with IPTG at 15°C for 16–18 hr. Cell lysis was performed in buffer A (20 mM Hepes/NaOH pH 7.5, 350 mM NaCl, 1 mM tris(2-carboxyethyl)phosphine (TCEP)), containing 10 mM imidazole. The lysate was bound to Ni-chelating sepharose beads in batch mode, and elution was performed in buffer A containing 400mM imidazole. Affinity tags were removed by 3C protease cleavage overnight at 4°C and applied to a S75 16/60 gel filtration column.

### In vitro hydroxylation assays

The thymidine hydroxylase activity assay was carried out in a reaction buffer containing: 50 mM HEPES/NaOH (pH7.6), 50 mM NaCl, 8 mM ascorbic acid, 4 mM 2-oxoglutarate, 1 mM Fe_2_SO_4_, 1 mM ADP, 20 μg mL^−1^ BSA and 0.5 mM DTT. The buffer was made anaerobic by degassing it with Argon for 1h at 4°C. The reaction was carried out at 37°C in a total volume of 50 μl, including 4 μM of the protein and 15μM of 14-mer double-stranded DNA (CAGCAGC**T**GCAACA). Upon completion of the reaction at indicated time points, samples were stored at −20°C for further prcessing.

### Sample preparation for mass spectrometry

Aliquots of 20 μL of the reaction mixtures were placed in 1.5 mL reaction tubes, and incubated at 95°C for 3 minutes followed by rapid cooling on ice, to denature the double stranded DNA to single stranded oligonucleotides. In each tube, 4 units of Nuclease P1 (Sigma-Aldrich, St. Louis, MO)) were added together with 100 μL digest buffer, containing 0.04 mM DFAM, 3.25 mM ammonium acetate pH 5.0 and 0.5 mM zinc chloride). The samples were incubated at 65 °C for 10 minutes to convert the oligonucleotides into single nucleotides. We then added 20 μL of Trizma base pH 8.5 and 4 units of alkaline phosphatase (Roche Life Science, Indianapolis, IN) and vigorously mixed for approximately 10 seconds. Samples were incubated at 37 °C (heating block) for 1 hour to allow for nucleotide to nucleoside conversion, after which we added 20 μL of 300 mM ammonium acetate pH 5.0 and evaporate to dryness at 40 °C in a TurboVap LV (Biotage, Uppsala, Sweden). Finally, we added 50 μL of 5 mM ammonium acetate in water – acetonitrile (2:98, v/v) and vigorously mixed for approximately 1 min.

### Measurement of hmU by mass spectrometry

Oligonucleotide HmU content was analysed as the released amount of 5′-hydroxymethyl-2′-deoxyuridine (HOMedU) after sample processing. A reference standard of HOMedU (Santa Cruz Biotechnology, Inc., Dallas, TX) was used to prepare calibration standards for HOMedU sample quantification.

For quantification, the HPLC-MS/MS system consisted of a QTRAP 5500 tandem mass spectrometer (Sciex, Framingham, MA, USA) coupled to an HPLC Acquity I Class pump (Waters, Milford, MA, USA). The HPLC system was equipped with a FTN I-Class autosampler and I-Class column oven (Waters). Data acquisition was performed using Analyst 1.6.2. software (Sciex).

The HPLC-MS/MS system was based on a previously developed method to quantify decitabine DNA incorporation^28^. This assay was modified to allow for HOMedU quantification in the positive electrospray ionization mode by using the following mass-to-charge ratio (m/z) transition: 257.0 → 124.0. The remaining settings of the method were unchanged.

### Preparation of J-DNA and JDBD:J-DNA and JBP1:J-DNA complex

J-DNA oligos^29^ were mixed with their complementary strand and annealed as previously described ^18^. Briefly, the hmC-containing oligonucleotide and the complementary strand were dissolved in water to a concentration of 100 μM and then heat-annealed. The double–stranded oligonucleotide was then glucosylated by the T4 Phage β-glucosyltransferase (T4-BGT) from New England BioLabs according to manufacturer’s instructions. To create the protein: J-DNA complexes, 1 mg ml^−1^ of JBP1 was mixed with J-DNA at 1:1.1 molar ratio and then concentrated with Amicon concentrators. The same procedure was used for making the JDBD:J-DNA sub-complexes. The sequences used in this study were 23-J-DNA TCGATT***<u>J</u>***GTTCATAGACTAATAC and J-15-DNA TAGAACCC***J***AACCAT.

#### SEC-SAXS data collection and analysis

Synchrotron X-ray data for all components were collected on a Pilatus 1M detector at the ESRF beamline BM29^30^. About 40 μl of each sample, at a concentration 3–10 mg ml^−1^ were loaded onto a Superose-6 column (Supplementary Table 1). The flow rate for SAXS data collection was 0.2 ml min^−1^ and a scattering profile was integrated every second. Frames for each dataset were selected based on the examination of the Size Exclusion profile together with the calculated R_g_ and D_max_ values. At least 20 frames for each dataset were selected, scaled and averaged using PRIMUS following the standard procedures (Supplementary Table 1). For the JDBD:23-J-DNA and JBP1:J-15-DNA, frames were analyzed with DATASW^31^.

### Model-independent analysis of SAXS data

SAXS data analysis was performed using the PRIMUS^32^ and ScÅtter^33^ software packages. The forward scattering I(0) was evaluated using the Guinier approximation (8) assuming the formula *I(q) = I(0)exp(-(qR_g_)^2^/3)* for a very small range of momentum transfer values (qR_g_< 1.3). Calculation of the pair distribution function and maximum distance D_max_ was performed using GNOM^34^. The R_g_ was estimated by Guinier approximation. The molecular mass was calculated using the Porod volume, and the Q_R_ method^33,35^. Ambiguity of all datasets was measured with AMBIMETER^36^. The useful range for each dataset was determined by SHANUM analysis^37^ prior to proceeding to *ab initio* modeling.

### *Ab initio* modeling using SAXS data

Molecular envelopes of *ab initio* created models, were made for all the components using DAMMIN^38^. Ten individual models were created for each component and averaging was performed using DAMAVER. Fitting of atomic resolution structures to molecular envelopes was performed using SUPCOMB^39^.

To resolve the relative positions of individual subunits of the JBP1 structure a volumetric analysis was performed using the program MONSA, an extension of the DAMMIN algorithm^38^, following an approach similar to the one described in^40^. MONSA allows *ab initio* modeling of macromolecular complexes by fitting simultaneously multiple experimental datasets. The search volume is defined as a sphere with radius equal to half the maximum dimension (D_max_) of the complex of study. A minimization algorithm based on simulated annealing fits the experimental datasets of each component (phase), while all components together should fit to the experimental dataset of the corresponding complex. In case of a multi-component complex, each different phase is assigned a different contrast (1 for protein, 2 for nucleic acid, 0 for solvent). In our approach we treat as a phase either different components (protein and DNA) but also the two distinct folding units that make JBP1 as we establish them in this work (ΔJDBD and JDBD). At least 20 individual runs for each complex (particle) were created. The online version of MONSA was used for all models generated in this study.

In case of a two-body modeling that consists of two components *X:y, X* denotes the component with the larger mass, and *y* the smaller mass. In our experiments, *X:y* would thus be JBP1:23-J-DNA or Δ-JDBD:JDBD. This modeling assumes that each component (phase) does not undergo conformational changes upon complex formation, or that the difference at that resolution is negligible.

Examination of the parameters stored in the log file, for each MONSA run show the fitting and the calculated *R_g_* values for each component from each individual MONSA run. To evaluate each MONSA modeling run we compared the *R_g_* derived from the experimental dataset for each component, with the calculated *R_g_* for each phase calculated for each MONSA model and stored in the log file. Runs with calculated *R_g_* values that differ significantly from the experimental-data derived *R_g_* were excluded from the analysis.

### Aligning models generated by MONSA

To compare MONSA models, we wanted to visualize the different position of each phase *y* relative to *X*, in a common reference framework. We therefore chose to first align all phases *X_i_* from each individual run. Then the transformation matrix of each X_i_ component was used to also transform the corresponding *y_i_* phase. The chosen procedure highlights the possible relative positions of *y_i_* relative to *X_i_*. This is preferable to aligning all complexes *X_i_:y_i_* to each other, which only yields the average relative position.

For that we developed a script in Python (available as supplementary information), that performs the following operations. DAMAVER is used to align all models of X_i_, the component that has the larger mass. DAMAVER superimposes all models to each other, averages them, selects the best model based on the normalized spatial discrepancy (NSD) metric as the reference model (X_ref_), and aligns all models to X_ref_. DAMAVER creates a new PDB file for each X_i_ in the new aligned position, also containing the transformation matrix (*T_i_*) that aligns each model (*X_i_*) with the reference (*X_ref_*). Then, the transformation matrix *T_i_* is used to transform every *y_i_* phase. Finally, DAMCLUST is used to cluster the transformed *y_i_* phases, effectively creating clusters that have the *y_i_* component in similar orientations. These clusters, define the conformational clusters of the to *X_i_:y_i_* complex.

Each cluster is presented as the average model of all *X_i_* components, and the average of the *y_i_* components for each cluster. Singleton clusters were excluded from our analysis. To evaluate the clustered models we examined their fit to the experimental data based on χ^2^ analysis.

To fit atomic resolution structures on molecular envelops generated by either DAMMIN or MONSA the program SUPCOMB^41^ was used.

To calculate the χ^2^ values of the hybrid dummy atoms model of Δ-JBP1 and the atomic model of JDBD:23-J-DNA we transformed the hybrid model using CRYSOL ^42^ and compared it to the experimental data of the complex.

### Structure similarity searches

Structure similarity searches were carried out by DALI searches against the whole PDB ^43^. DALI returned new hits compared to previous searches^44^. We then inspected the top hits manually. The recognition helix, the supporting helix, as well as two of the other helices and the connectivity we have described for the helical bouquet fold of JDBD^44^ were present in the top 1 (self), 2, 3, and 5 hits. Hit nr. 5 (Z-score 5.2, RMSD 3.0, 10% identity over 84 aligned residues) is the motility gene receptor MogR that we have previously identified as a JDBD structural homolog. Hit nr. 2 (Z-score 6.7, RMSD 3.0, 10% identity over 97 aligned residues) is AcrF3, a protein encoded by gene 35 from phage JBD5 that has been reported to specifically inhibit the Cas3 protein of *P. aeruginosa* strain UCBPP-PA14 (PaCas3) and to counteract the type I–F CRISPR–Cas system^45^. Hit nr. 3 (Z-score 6.0, RMSD 4.0, 11% identity over 94 aligned residues) is a chromodomain helicase DNA binding protein. Hit 4 has the lowest sequence identity (6%), the recognition helix is missing, and is a potassium channel with no functional homology. Hits in position 5 and below were too distant to consider, as judged by Z-scores (4.8 and below), RMSD (4.5 and above) and had no functional similarity (DNA binding).

## Supplementary Information

**Supplemental Table 1.**
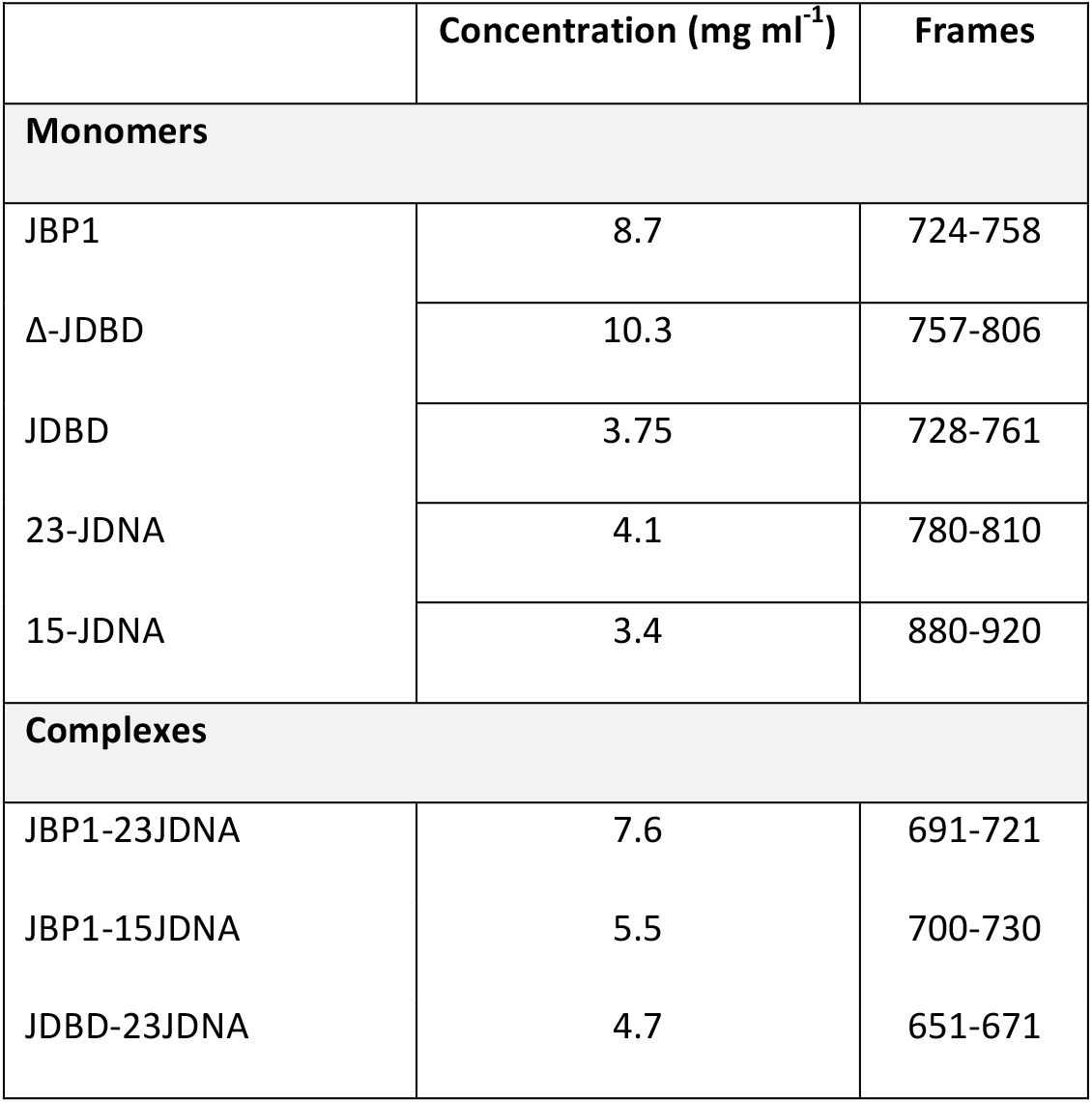

**Supplementary Figure 1.**
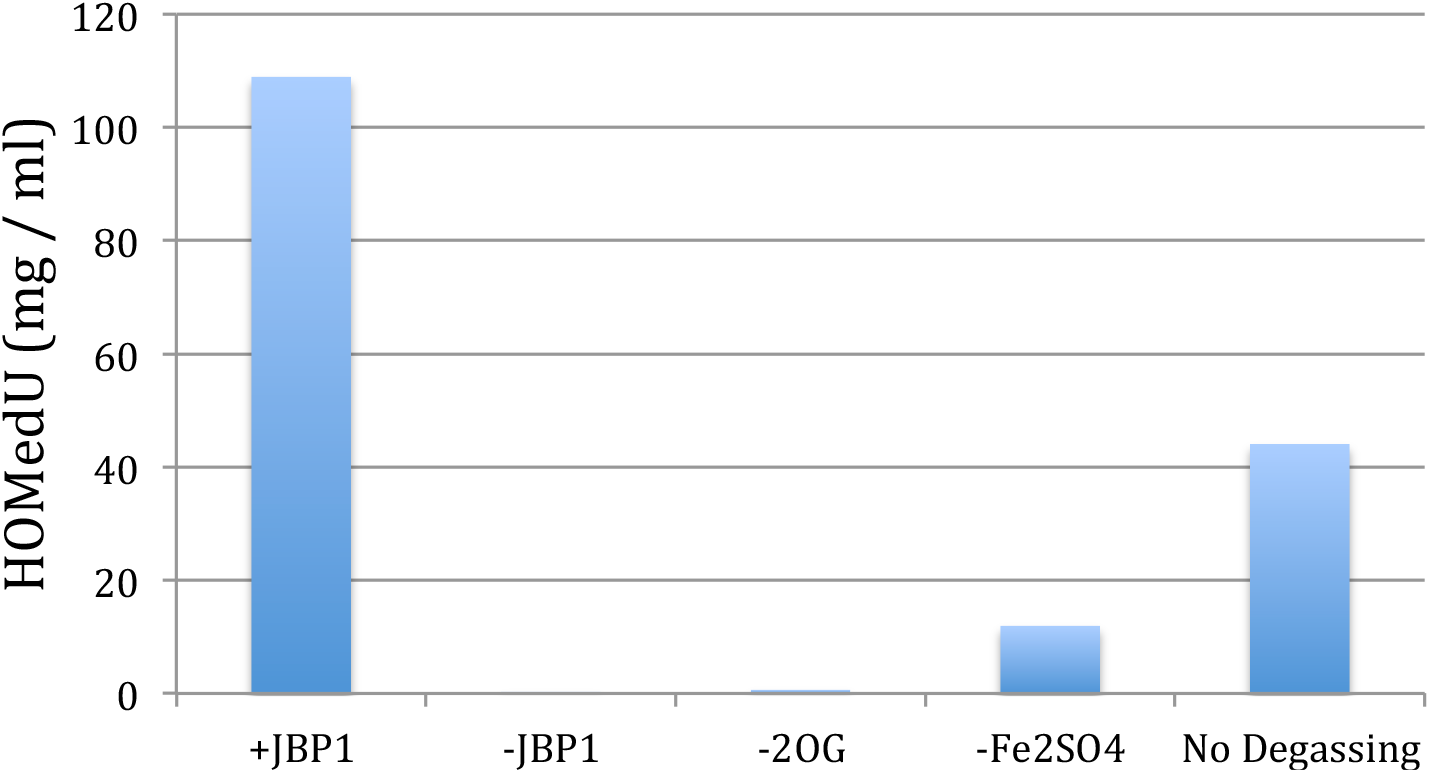
Amount of hmU produced by JBP1 under the conditions we describe in methods, and related controls without protein, co-factor, iron, and without degassing.

**Supplementary Figure 2.**
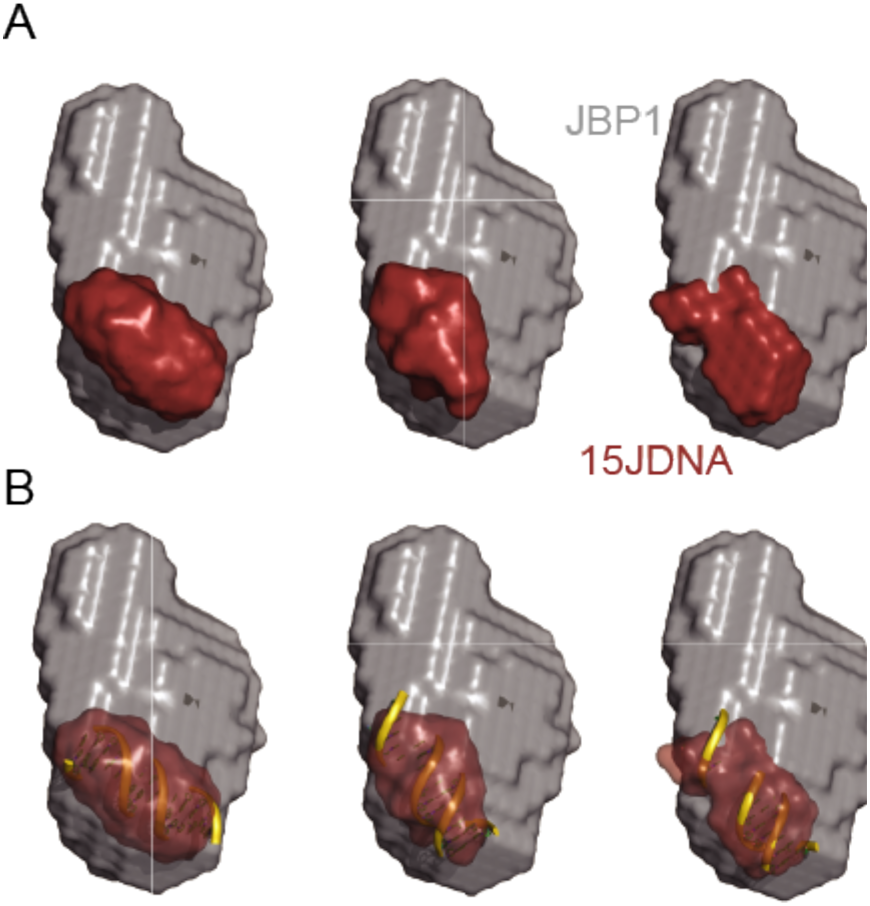
Model of JBP1:J-15-DNA. **(A)** Clusters of all JBP1:J-15-DNA MONSA models, with the χ^2^ fitting of the calculated intensities to the experimental data. **(B)** The J-15-DNA is superimposed on the molecular envelope generated by MONSA. Superposition was performed using SUPCOMB from the ATSAS suite

**Supplementary Figure 3.**
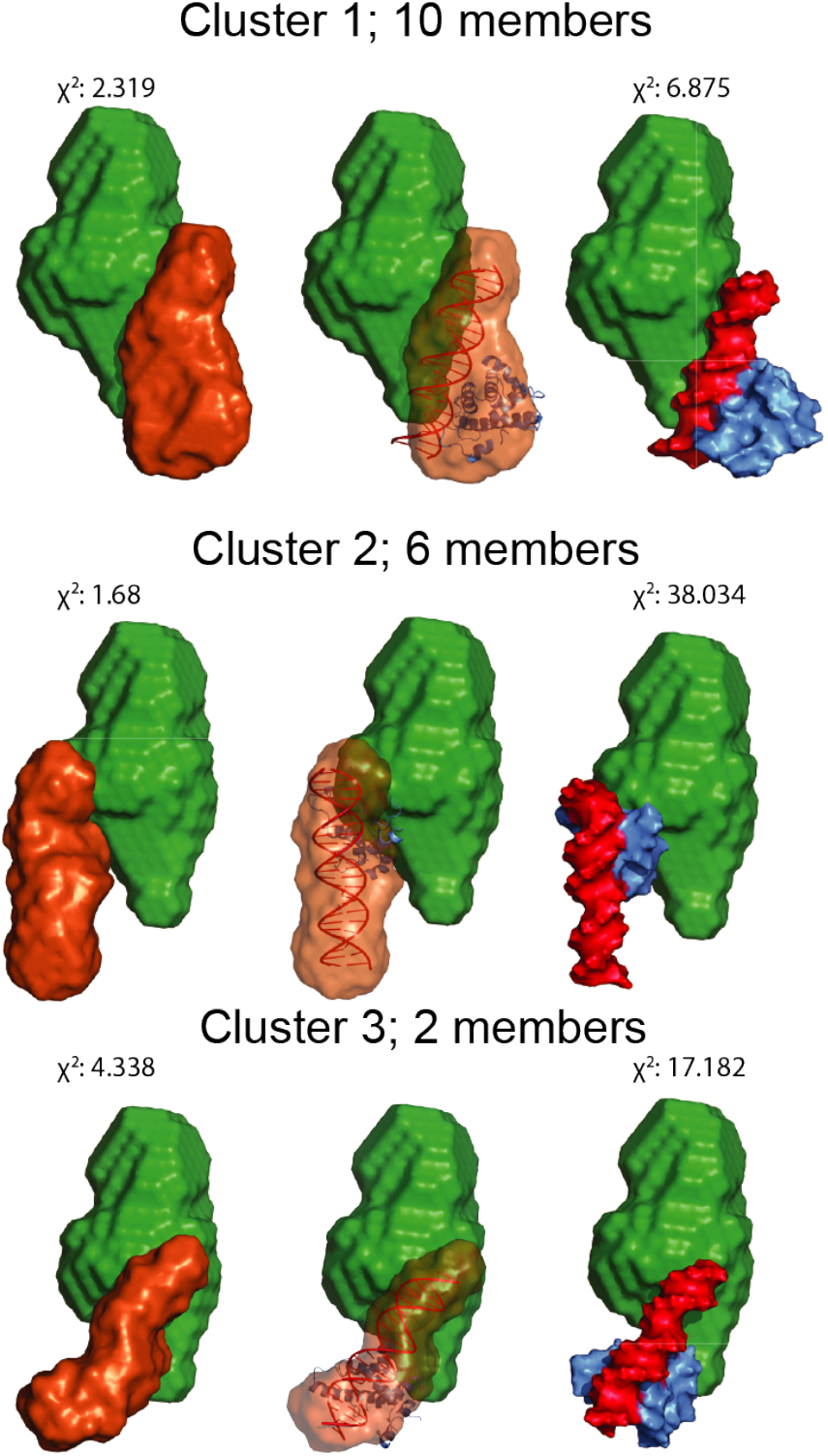
All MONSA model clusters of JBP1 in presence of JDNA. From left to right, MONSA models of the relative positions of Δ-JDBD and JDBD:J-23-DNA sub-complex, a pseudo-atomic model of JDBD:J-23-DNA fitted in the molecular envelope of the generated model; and a surface representation of all components. Figures were made using Pymol.

